# Inhibition of host Lactate dehydrogenase A by a small-molecule limits *Mycobacterium tuberculosis* growth and potentiates bactericidal activity of isoniazid

**DOI:** 10.1101/626002

**Authors:** Gopinath Krishnamoorthy, Peggy Kaiser, Ulrike Abu Abed, January Weiner, Pedro Moura-Alves, Volker Brinkmann, Stefan H. E. Kaufmann

## Abstract

Lactate dehydrogenase A (LDHA) mediates interconversion of pyruvate and lactate. Increased lactate turnover is shared by malignant and immune cells. Hypoxic lung granuloma in *Mycobacterium tuberculosis*-infected animals present elevated levels of *Ldha* and lactate. Such alteration in metabolic milieu could influence the outcome of interactions between *M. tuberculosis* and its infected immune cells. Given the central role of LDHA for tumorigenicity, targeting lactate metabolism is a promising approach for cancer therapy. Here, we sought to determine the importance of LDHA for Tuberculosis (TB) disease progression and its potential as a host-directed therapeutic target. To this end, we administered FX11, a small-molecule NADH-competitive LDHA inhibitor, to *M. tuberculosis* infected C57BL/6J mice and Nos2^−/−^ mice with hypoxic necrotizing lung TB lesions mimicking human pathology more closely. FX11 did not inhibit *M. tuberculosis* growth in aerobic/hypoxic liquid culture, but modestly reduced the pulmonary bacterial burden in C57BL/6J mice. Intriguingly, FX11 administration limited *M. tuberculosis* replication and onset of necrotic lesions in Nos2^−/−^ mice. In this model, Isoniazid (INH) monotherapy has been known to exhibit biphasic killing kinetics owing to the probable selection of an INH-tolerant subpopulation. This adverse effect was corrected by adjunct FX11 treatment and augmented the INH-derived bactericidal effect against *M. tuberculosis*. Our findings therefore support LDHA as a potential target for host-directed adjunctive TB therapy and encourage further investigations into the underlying mechanism.

**IMPORTANCE:** Tuberculosis (TB) continues to be a global health threat of critical dimension. Standard TB drug treatment is prolonged and cumbersome. Inappropriate treatment or non-compliance results in emergence of drug-resistant *Mycobacterium tuberculosis* strains (MDR-TB) that render current treatment options ineffective. Targeting the host immune system as adjunct therapy to augment bacterial clearance is attractive as it is also expected to be effective against MDR-TB. Here, we provide evidence that pharmaceutical blockade of host lactate dehydrogenase A (LDHA) by a small-molecule limits *M. tuberculosis* growth and reduces pathology. Notably, LDHA inhibition potentiates the effect of Isoniazid, a first-line anti-TB drug. Hence, its implications of our findings for short-term TB treatment are profound. In sum, our findings establish murine LDHA as a potential target for host-directed TB therapy.

## INTRODUCTION

Tuberculosis (TB) is the leading cause of mortality from an infectious agent globally (1) and its treatment includes six-month long therapy with combinations of drugs. Development of newer drugs with superior efficacy and safety is urgently required to shorten the treatment duration as well as to manage drug-resistant TB effectively. Pathogen-targeted treatment is the preferred choice, however, host-directed approaches are being increasingly recognized for adjunct therapy to reduce pathogen load and ameliorate exacerbated organ damage during TB granuloma progression (2, 3). Radiotracer imaging of *M. tuberculosis*-infected lungs has revealed heterogeneity – in size, metabolism, and infection – within and between granulomas in a single host (4, 5). In general, the significance of metabolic processes on immune functions is increasingly accepted (6–8). Heterogeneous responses in granuloma, therefore, could partly be attributed to metabolic state(s)/energy phenotype(s) of different immune cells (e.g., macrophages, neutrophils, lymphocytes) that are influenced by their microenvironment and local infection dynamics. Understanding of pathogen-induced immunometabolic dysregulation in granuloma can provide insights into the vital pathways in the infected host and thereby reveal novel therapeutic target candidates.

Untargeted metabolite analysis has identified elevated levels of lactate in necrotic granuloma of *M. tuberculosis*-infected guinea pigs (9). Generation of lactate from pyruvate, a terminal glycolytic step, is catalyzed by lactate dehydrogenase A (LDHA), whose functions depend on hypoxia-inducible factors (HIFs) (10). Both LDHA and HIF1-α transcripts have been found to be significantly induced in *M. tuberculosis*-infected mouse lungs (11, 12), and the essential function of HIF1-α in controlling TB progression has already been recognized (10). Although metabolic phenotypes of malignant and immune cells show some critical differences, they present many similarities (13). In most cancer cells, aerobic glycolysis (Warburg effect) or hypoxia adaptation requires LDHA, and its inactivation using the NADH competitive inhibitor, FX11 (3-dihydroxy-6-methyl-7-(phenylmethyl)-4-propylnaphthalene-1-carboxylic acid; PubChem CID: 10498042), has been shown to regress lymphoma and pancreatic cancer (14). In this report, we interrogate whether FX11-mediated LDHA inhibition could result in host-beneficial and pathogen-detrimental outcome in murine TB models and its relevance to host-directed therapy.

## FINDINGS

FX11 affects bioenergetics and glycolysis in human Panc (P) 493 B-lymphoid cells (14). Here, we assessed the FX11-induced response in interferon-gamma (IFN-γ) stimulated but uninfected murine bone marrow derived macrophages (BMDMs) **(Methods in Text S1)**. FX11 addition increased the oxygen consumption rate (OCR), but decreased the respiratory capacity, membrane potential, and ATP synthesis in a concentration-dependent manner (Fig. 1A and B**; Text S2**). Essentially, FX11 (at 14.3 µM) uncoupled the mitochondrial respiratory chain and phosphorylation system. Likewise, FX11-mediated LDHA inhibition increased the extracellular acidification rate (ECAR) (implying increased glycolysis) but depleted the cellular glycolytic reserve (Fig. 1C and D). Such FX11-dependent glycolytic induction could be argued, in part, as a measure to compensate the reduced mitochondrial energy generation. Nevertheless, these observations establish that FX11-mediated LDHA inhibition profoundly affects bioenergetics and glycolysis in BMDMs. Intriguingly, recent studies have demonstrated that energy-flux changes in macrophages depend on viability and virulence of *M. tuberculosis* (15, 16). Upon virulent *M. tuberculosis* infection, human monocyte-derived macrophages shift their energy generation to mitochondrial fatty acid oxidation with concomitant decrease in glycolysis (16). We interrogated whether FX11-mediated impairment of respiratory/glycolytic function directly affects the intramacrophage *M. tuberculosis* survival. Because high concentration of FX11 affected viability of BMDMs, we tested 1.43 μM concentration and the bacterial survival remained identical between untreated and FX11-treated conditions (Fig. 1E).

Although FX11 is an analog of anti-bacterial gossypol (17), we found that FX11 is non-toxic to *M. tuberculosis* under the tested conditions. The aerobic growth of *M. tuberculosis* in glycerol or sodium L-lactate was comparable between FX11-treated and untreated growth control (Fig. 1F and G). Similarly, fluorescence measurement of green fluorescent protein expressing *M. tuberculosis*, as a function of growth, under 1% O_2_ hypoxia revealed that FX11 did not affect the bacterial viability, albeit a minor decrement in fluorescence was noted (Fig. S1A and B). Moreover, development of a pale brownish color in FX11-supplemented hypoxic culture only was noted suggesting that this small-molecule is differentially metabolized under such condition. Finally, the respiratory functions in *M. tuberculosis* also remained unperturbed when FX11 was added (Fig. 1H). We conclude that the bioenergetics effects of FX11 are highly host-specific.

**FIG 1.**
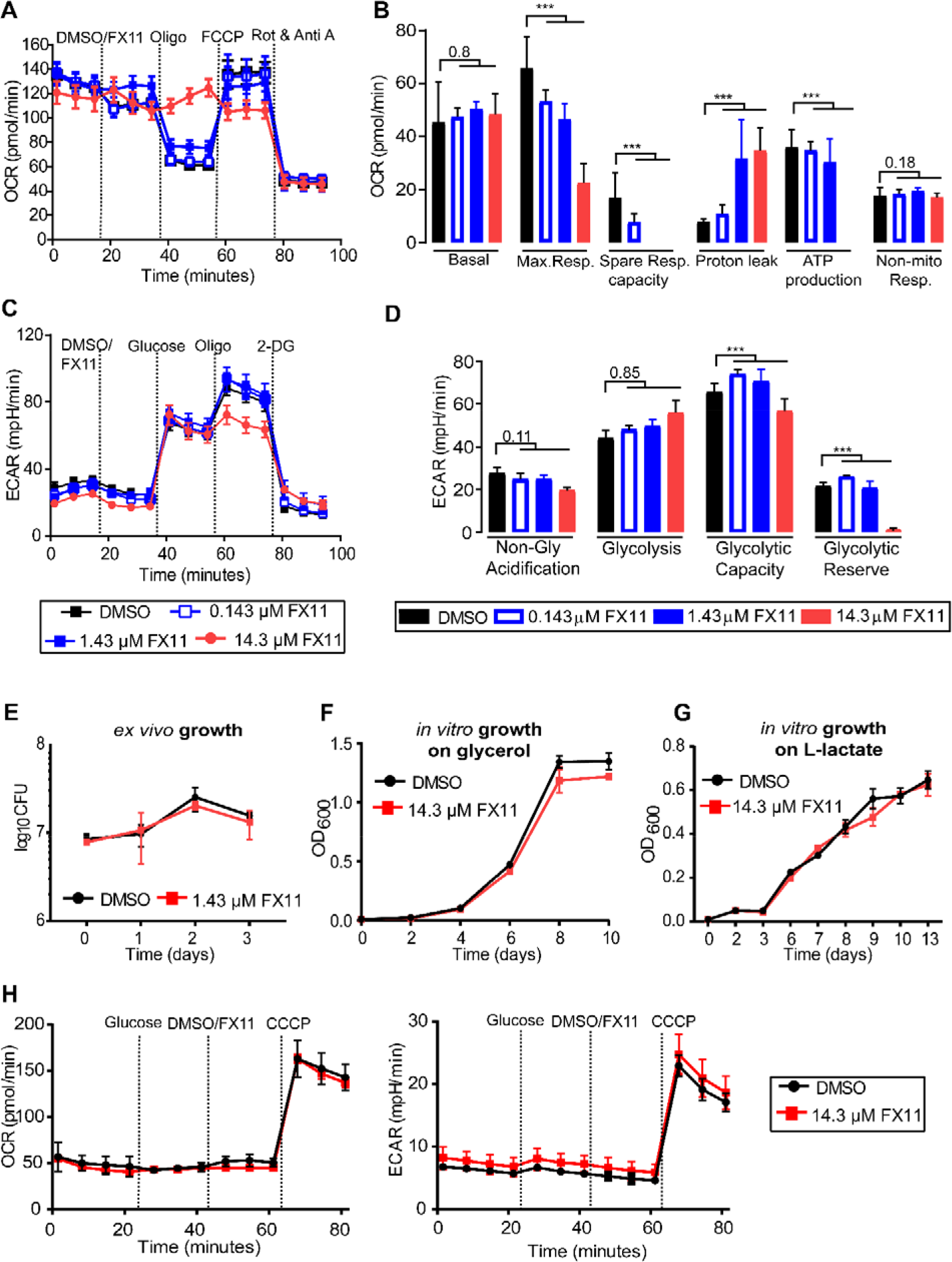
FX11-induced metabolic changes are highly host-specific. FX11 alters the **(A-B)** respiratory profile and parameters, **(C)** glycolytic parameters, and **(D)** glycolytic proton efflux rate (PER) of IFN-γ-stimulated murine bone marrow-derived macrophages (BMDMs) in a concentration-dependent manner. Wells with DMSO served as control. Different mitochondrial and glycolytic modulators were sequentially injected and cellular responses (OCR and ECAR values) were measured using Seahorse XF analyzer. Data represent three independent experiments. A regression model was performed to determine the dose-response effect of FX11 on BMDM metabolic parameters using the pooled data from three independent experiments *, P >0.001; ***, P >0.0001 (*see* **Text S2**). **(E)** IFN-γ-stimulated BMDMs infected with *M. tuberculosis* H37Rv at multiplicity of infection 1:5, with FX11 effect determined by enumerating viable bacterial counts. Effect of FX11 on *M. tuberculosis* growth in liquid medium containing **(F)** 0.2% v/v glycerol or **(G)** 10 mM sodium L-lactate as the sole carbon source. **(H)** Effect of FX11 *M. tuberculosis* respiratory function (OCR and ECAR values) measured by Seahorse XFp extracellular flux analyzer.

Subsequently, the effect of FX11 was evaluated in two murine TB models. In a first experiment, C57BL/6J mice were aerosol-infected with 100 CFU of *M. tuberculosis* H37Rv. At 4 weeks post-infection, mice received either FX11 (2 mg/kg) or 2% (final concentration) dimethyl sulfoxide (DMSO) as placebo by oral gavage (6 days/week) for further 4 weeks. Post-treatment effect was monitored at 2 and 4 weeks by enumerating CFU from excised lungs and spleens of euthanized animals. FX11 administration resulted in approximately 0.5 log_10_ reduction in pulmonary *M. tuberculosis* counts (Fig. 2A; Fig. S2A) with less apparent effect on splenic CFU. Administered dose of FX11 is similar to those in a previous study (14), and further dose increment is restricted due to poor compound solubility. Furthermore, complete inhibition of LDHA could result in adverse events as it is essential for cellular homeostasis.

TB granulomas in C57BL/6J mice rarely progress into necrosis, whereas Nos2^−/−^ mice present hypoxic necrotizing lung lesions that are characteristics hallmarks of human TB (18–20). Therefore, in a second experiment, the effect of FX11 (2 mg/kg), either individually or in combination with isoniazid (INH, 25 mg/kg), was evaluated in Nos2^−/−^ mice (Fig. 2B). Efficacy was determined by assessing histopathology and bacterial viability. FX11 administration was apparently well-tolerated because treated animals showed no increased distress or weight loss (Fig. S2B). As previously observed (20), onset of hypoxic and necrotic lesions became apparent at day 56 (at treatment start). Although the number and size of lesions were comparable, further development of necrotic lesions were ceased in the FX11-treated group (Fig. 2C,D; Fig. S2C,D). Likewise, 2 or 4 weeks of FX11 administration limited further bacterial growth in lungs and spleens. Immunofluorescence staining of paraffin-embedded lung sections revealed that LDHA expression co-localized with hypoxic lesions (Fig. 2C; Fig. S3). Nonetheless, FX11 administration had no apparent impact on LDHA immunofluorescence which is probably due to non-inhibitory effects of FX11 on transcription/translation. Moreover, enzymatic quantification of lactate from the excised lung tissues presented erroneous and irreproducible data (data not shown). Thus, no concrete evidence could be presented to corroborate the *in vivo* inhibitory effect of FX11 on LDHA.

Necrotic lesions in the Nos2^−/−^ model have been correlated with the evolution of slow/non-growing INH-tolerant subpopulation (20). Accordingly, we interrogated whether FX11-mediated inhibition of progression to necrotic granuloma potentiates INH efficacy possibly by preventing the emergence of the drug-tolerant population. Indeed, the combination of FX11 and INH resulted in superior efficacy, and there was no further cessation of bactericidal activity of INH, when compared with monotherapy (Fig. 2B). While this observation requires further validation in other experimental models, it has immense implications for shortening TB treatment and minimizing the risk of emergence of drug resistance in *M. tuberculosis.*

**FIG 2.**
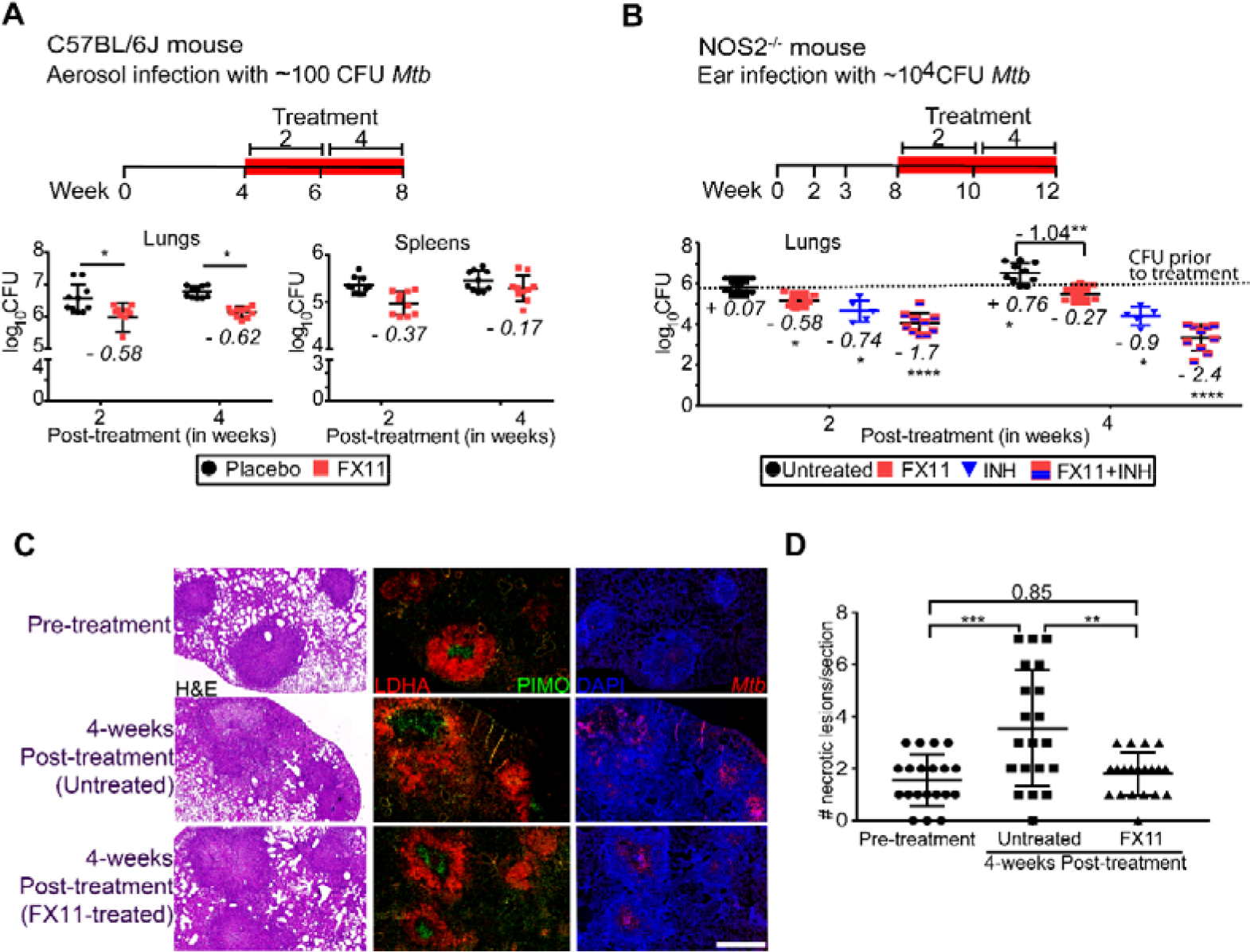
Evaluation of FX11 effects against *M. tuberculosis* in mouse models. **(A)** Schematic representation of experimental design (treatment duration is highlighted in red). Effect of FX11 (2 mg/kg) on bacterial burden in C57BL/6J mice aerosol infected with 100 CFU *M. tuberculosis*. Datasets presented are from 2 independent experiments (n = 10). Values shown are means±standard deviation (SD). Italicized numerical value (in negative) represents reduction in log_10_CFU in the treated group, when compared with the placebo control group. Statistical significance was evaluated by using an unpaired Student *t* test. *p<0.05. **(B)** Effect of FX11 (2 mg/kg) as monotherapy or in combination with INH (25 mg/kg) in Nos2^−/−^ mice with hypoxic necrotizing lung lesions (20). The TNF-α response was neutralized at 2 and 3 weeks of post-infection. Drugs were administered after onset of central necrosis and hypoxia in lung lesions at day 56. Untreated or INH-treated (n = 5) groups were used for comparisons. Lung CFU data (means±SD) from two independent experiments (n = 9–10) are shown. Italicized numerical value represents log_10_CFU differences (an increase is indicated as positive value and a decrease is in negative integer) of the specified group, when compared with the control group prior to drug treatment (i.e. day 56, indicated in dotted line). Pooled data from two independent experiments were analyzed using nonparametric Mann-Whitney test (data that did not pass the Shapiro-Wilk normality test). Statistical significance as compared to the group prior to drug treatment, *p<0.05, **p<0.01, ****p<0.0001. **(C)** Hematoxylin and eosin (H&E) staining and immunofluorescence detection of *M. tuberculosis* or hypoxia marker pimonidazole (PIMO) and LDHA. Magnified images show the co-localized staining of LDHA and PIMO in lung lesion. Scale bar represents 1 mm. Micrographs of a stained section of whole left lung lobe are presented in Fig. S3. **(D)** Total number (means±SD) of necrotizing lesions present in Nos2^−/−^ mice that were either untreated or FX11-treated. Data were analyzed using two-way ANOVA with multicomparison and Tukey’s post-test. Statistical significance as compared to the control group prior to drug treatment, n=5, **p<0.01, ***p<0.001.

Genetic ablation of LDHA in T cells has been found to protect mice from IFN-γ-mediated lethal pathology of autoimmune responses (21). Similarly, lactate accumulation has been indicated to severely impair IFN-γ-dependent tumor immunosurveillance (22). It is a well-established paradigm that IFN-γ has a central role in macrophage activation and tissue-protection in TB (23, 24). Besides, HIF-1α is not only a transcriptional regulator of LDHA, but also coordinates IFN-γ-dependent adaptive immunity to *M. tuberculosis* (10). It has been reported that IL-17 limits HIF1α expression (and lactate accumulation) and hypoxic necrotic granuloma development in C3HeB/FeJ mice infected with an *M. tuberculosis* clinical isolate (12). Thus, LDHA inhibition resulting in heightened IL-17 activity and/or reduced IFN-γ-dependent exacerbated inflammation could explain the FX11-limited necrotic granuloma progression in the Nos2^−/−^ mouse model. However, the cause of reduction in *M. tuberculosis* burden upon FX11 treatment is difficult to explain. FX11-mediated LDHA inhibition perhaps alters the balance of pro- and anti-inflammatory cytokines thereby contributing to *M. tuberculosis* clearance (25). Observed FX11effects can also be linked to factors other than LDHA inhibition. E.g., reactive catechol moiety of FX11 or its drug-intermediates (under oxygen limiting conditions) could cause off-target effects. FX11 has been shown to induce oxidative stress (14), which could restrict bacterial growth and augment INH efficacy against *M. tuberculosis*. Finally, FX11 administration could deprive *M. tuberculosis* from utilizing host-derived lactate for energy generation (26). Therefore, in depth analysis of mechanism underlying LDHA inhibition and pathogen clearance is warranted as it is a promising host-directed therapy approach in adjunct to canonical drug treatment.

## ACKNOWLEDGEMENTS

Animal protocols were approved by Landesamt für Gesundheit und Soziales, Berlin, Germany. Experiments were conducted in accordance with the European directive 2010/63/EU on Care, Welfare and Treatment of Animals.

We thank Manuela Primke, Ines Neumann, Jens Otto, Uwe Klemm, and Gesa Rausch for their help in mouse breeding and maintenance; Marion Klemm, Manuela Stäber and Dagmar Oberbeck-Mueller for technical assistance. We gratefully acknowledge the partial financial support (to S.H.E.K) from “PreDiCT-TB” and the intramural funding of Max Planck Society to S.H.E.K.

## TEXT S1. Supplementary material and methods

### Bacterial strains

*M. tuberculosis* H37Rv (American Type Culture Collection, #27294) or its derivative expressing pGFPHYG2 replicative plasmid (kind gift from Lalita Ramakrishnan; Addgene# 30173) was grown in Middlebrook 7H9 broth (Becton Dickinson) supplemented with albumin-dextrose-catalase enrichment (Becton Dickinson), 0.2% glycerol, 0.05% Tween 80 or on Middlebrook 7H11 agar (Becton Dickinson) containing 10% v/v oleic acid-albumin-dextrose-catalase enrichment (Becton Dickinson) and 0.2% glycerol. 10 mg of FX11 (Merck Millipore) was dissolved in 1 mL of dimethyl sulfoxide (DMSO).

### Growth assay

Bacterial growth (with 5% DMSO or 14.3 μM FX11) was assessed in minimal medium (0.5 g/liter asparagine, 1 g/liter KH_2_PO_4_, 2.5 g/liter Na_2_HPO_4_, 50 mg/liter ferric ammonium citrate, 0.5 g/liter magnesium sulfate, 0.5 mg/liter calcium chloride, and 0.1 mg/liter zinc sulfate) containing either 0.2% glycerol (vol/vol), or 0.5% glucose (wt/vol), 0.01% cholesterol (wt/vol), 10mM sodium L-lactate. Cell densities (OD) were measured at 600 nm by using a cell density meter (BioChrome Biowave). Infection stocks were prepared from mid-log phase *M. tuberculosis* cultures. For CFU determinations, serial dilutions were performed in PBS/0.05% Tween 80 and plated onto Middlebrook 7H11 agar. Plates were incubated at 37 °C for 4–5 weeks prior to CFU counting. For fluorescent-based-measurement, black, optical-bottom, 96-well microplates was used and fluorescence measured with a GloMax^®^ Microplate Multimode Reader using “Blue” filter (Excitation: 490 nm, Emission: 510–570 nm).

### Drugs, formulations and administration

FX11 (Merck Millipore) or INH (Sigma) were formulated in 0.4% methylcellulose. The final concentration of DMSO did not exceed 2%. Drug formulations were prepared every week and stored at 4 °C. Drugs were administered by oral gavage (0.2 ml) on 6 days per week.

### Ethical statement

All animal studies have been ethically reviewed and approved by the State Office for Health and Social Services, Berlin, Germany. Experimental procedures were carried out in accordance with the European directive 2010/63/EU on Care, Welfare and Treatment of Animals.

### Animal experiments

Female C5BL/6J and C57BL/6J Nos2^−/−^ mice were bred in-house and maintained under specific pathogen-free conditions. Six- to eight-week-old C5BL/6J mice were aerosol infected with 100 CFU *M. tuberculosis* H37Rv. C5BL/6J Nos2^−/−^mice were infected as previously reported (2). In brief, six- to eight-week-old female C5BL/6J Nos2^−/−^mice were anesthetized (ketamine 65mg/kg, acepromazine 2 mg/kg, xylazine 11 mg/kg) and infected with 1,000 CFU of *M. tuberculosis* in 20 μl PBS given into the ear dermis. At 14 and 21 days post-infection each mouse received 0.5mg of monoclonal anti-tumour necrosis factor alpha antibody (purified from MP6-XT22 cultures) by intraperitoneal (i.p.) injection. Two hours before euthanasia animals received 60mg/kg pimonidazole hydrochloride (Hypoxyprobe™-1, Burlington, MA, USA) i.p. to allow for detection of hypoxic regions in organ sections.

### Staining procedures and histopathology

The left lung lobe of mice was removed aseptically and post-fixed in 4% paraformaldehyde for 16–20 h at room temperature. The tissue was then dehydrated and paraffin-embedded (60 °C) using a Leica TP 1020 tissue processor. Paraffin blocks were cut at 2-3 μm, sections were mounted and dried on Superfrost Plus slides (Thermo Scientific) avoiding temperatures above 37 °C. After dewaxing and rehydration, sections were subjected to haematoxylin and eosin (H&E) staining, or fluorescence staining to detect LDHA expression, pimonidazole and *M. tuberculosis* in tissues. Sections were stained with hematoxylin/eosin using standard protocols. Central necrosis of lesions was defined as a lighter pink region indicating tissue consolidation surrounded by granulomatous inflammatory infiltrates. Researcher blinded to the study groups scored at least 4 individual stained sections of each organ in study groups of five mice per time point.

For immunostaining, sections were incubated in one of the heat-induced epitope retrieval (HIER) buffers (pH 6, citrate) for 20 min at 96 °C in a steam cooker (Braun). After antigen retrieval, sections were left in the same HIER buffer at room temperature to cool below 30 °C. Sections were further rinsed three times with deionized water and once with Tris-buffered saline (TBS, Pierce Protein-Free Blocking Buffer (pH7.4)). Subsequently sections were permeabilized for 5 min with 0.5% Triton X100 in TBS at room temperature, followed by three rinsing steps with TBS. Sections were surrounded with PAP pen and treated with TBS blocking buffer for 30 min to prevent non-specific binding. Primary antibodies were diluted in TBS blocking buffer and incubated on the sections over night at room temperature.

Following antibodies were used for immunostaining: Anti-*Mycobacterium tuberculosis* antibody (Abcam, ab905), Anti-LDHA antibody (Abcam ab101562 LOT GR176934), anti-pimonidazole (PIMO) primary antibody is FITC-conjugated (included in kit), secondary detection of PIMO is carried out using goat anti-FITC (Abcam ab19224 LOT GR175456-35) followed by donkey anti-goat Alexa 488 (Dianova 705-546-147). The following antigen retrieval solutions were used: R-Universal Buffer pH7, 10× (Aptum APO 0530500), Target Retrieval Solution pH9 10mM Tris (TRS) 10× (Dako S236784), and Target Retrieval Solution pH6 10mM Citrate 10× (Dako S236984-3). Dilution and blocking buffer were TBS supplemented with 1% BSA/2% donkey NS/5% cold water fish gelatin/0.05% Tween 20/0.05%Triton X100.

Fluorescence images were recorded using a Leica SP8 confocal or a Leica DMR widefield microscope (equipped with bandpass filter blocks and a Jenoptik ProgRes MF USB camera). Complete tissue sections were digitized using a ZEISS Axioscan Z1 slide scanner.

### Bacterial enumeration from lungs and spleens

Mice were euthanized at dedicated time points and superior, middle inferior and post-caval lobes were removed and homogenized in 1ml PBS/0.05% Tween 80. Serial dilutions of organ homogenates were plated onto Middlebrook 7H11 agar and in addition on agar supplemented with 0.4% activated charcoal for all time points during chemotherapy. Plates showing higher CFU counts were used for data analysis.

### Isolation of bone marrow derived macrophages (BMDMs)

BMDMs was obtained from tibia and femur bones and maintained in Dulbecco’s Modified Eagle Medium containing 20% L929-cell supernatant, 10% heat-inactivated FCS, 5% heat-inactivated HS, 2 mM glutamine. Differentiated resting cells and cells pretreated with recombinant mouse IFN-γ (100 U/ml; Strathmann Biotech AG) were infected with *M. tuberculosis* H37Rv at MOI 1:5.

### Extracellular flux analysis

Seahorse Xfp extracellular flux analyzer (Agilent, Santa Clara, CA) was used to measure oxygen consumption (OCR) of *M. tubeculosis* cells as described earlier (3,4) and Seahorse XF96 extracellular flux analyzer (Agilent, Santa Clara, CA) was used to measure oxygen consumption (OCR) and extracellular acidification rates (ECAR) of murine BMDM cells as per manufactures recommendation. Cells were seeded into the XF96 cell culture plate at cell densities of 70000 cells/well and rested for 24 h. Subsequently cells were stimulated with IFN-γ (100 U/ml) for further 24 h at 37 °C /7% CO_2_.

Mitochondrial respiration assay (Seahorse XF cell mito stress test) and glycolytic function assay (Seahorse XF Glycolysis Stress Test) were performed according to the manufacture’s recommendation. Data analysis was carried out using The Wave Desktop 2.6 Software (available at https://www.agilent.com/en/products/cell-analysis/software-download-for-wave-desktop) and the XF Report Generators for calculation of the parameters from the respective assays.

Assay principle, design, and equations to calculate each of the parameters is schematically illustrated below using the representative data obtained in this study. A more detailed account of these assays can be accessed from the manufacturer’s web resources.

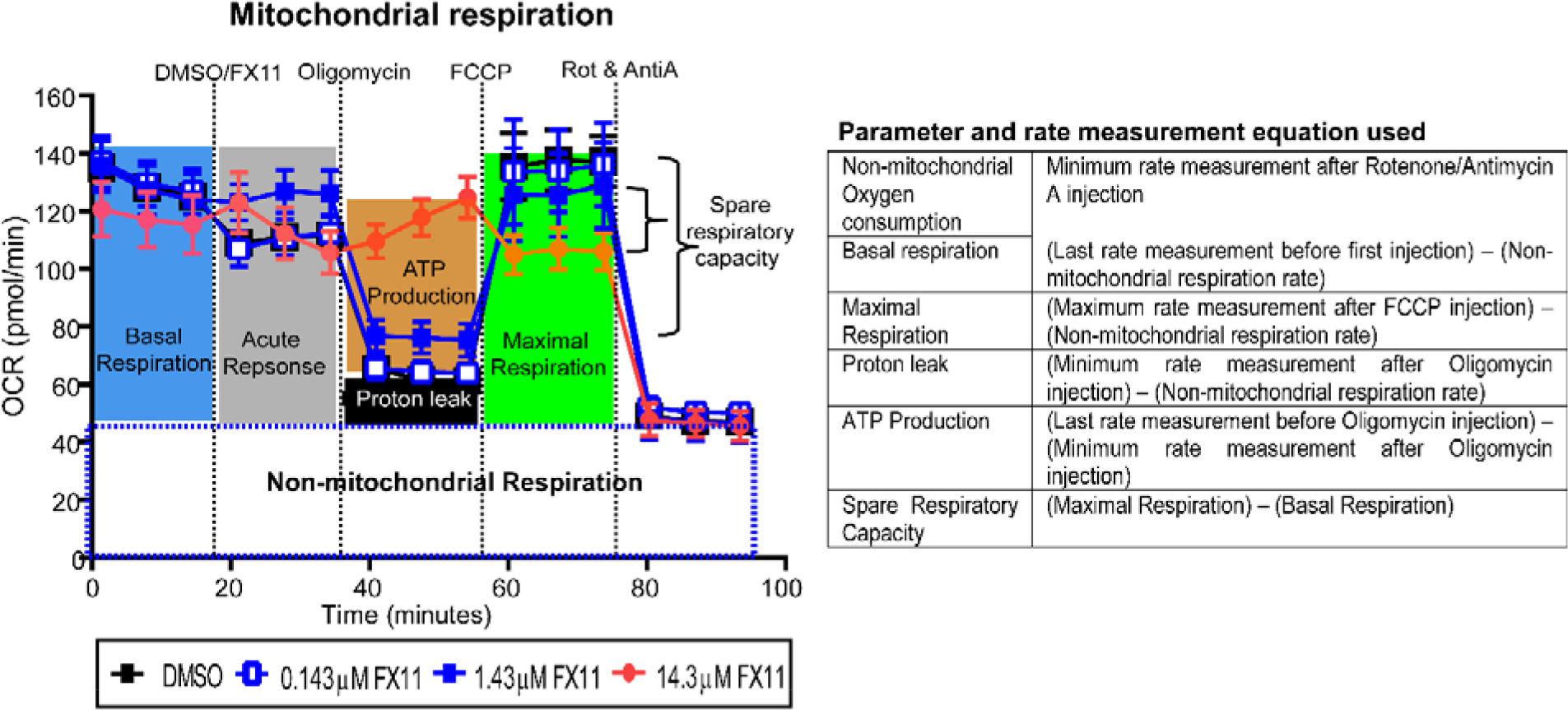

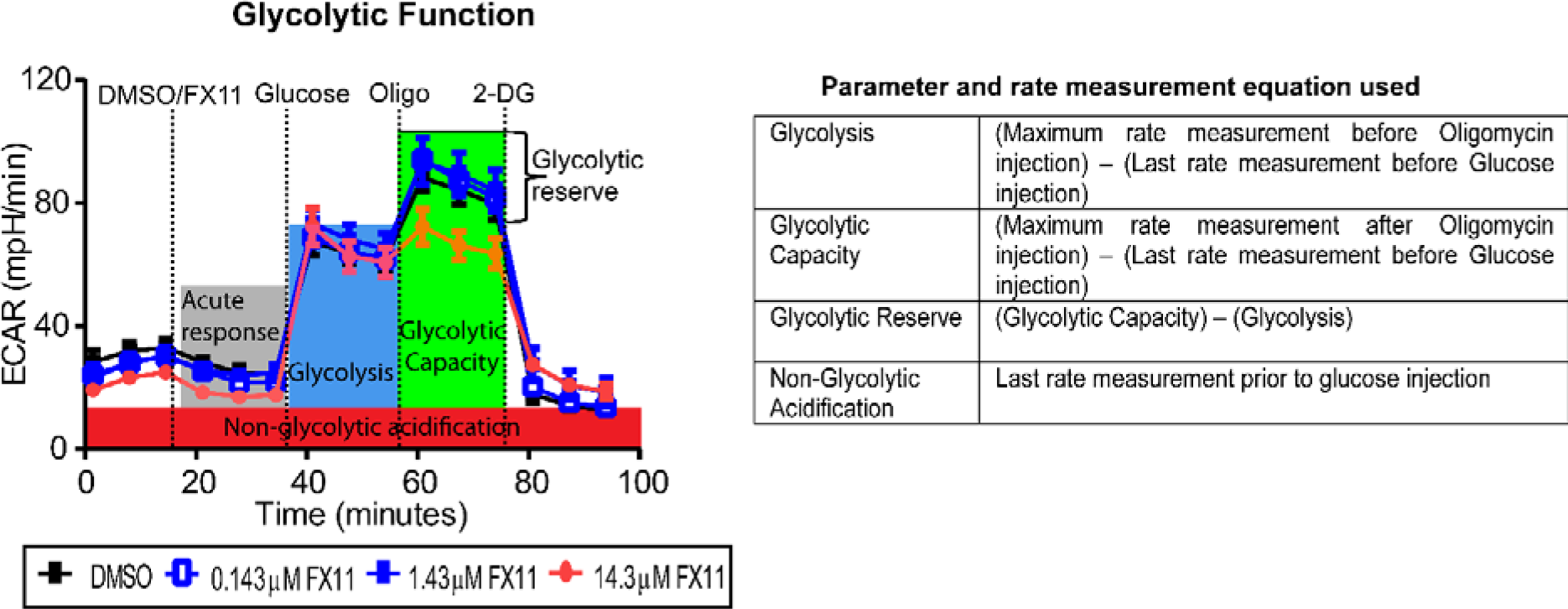

### Assay type, injection sequence of modulators used in this study

#### Mitochondrial respiration assay

|  | **Mitochondrial respiration** |  |  |
| --- | --- | --- | --- |
|  | **Compound** | **Function** | **Effect on OCR** |
| **Basal respiration** | Not applicable | - | To monitor the cellular energetic demand under baseline conditions. |
| **Injection 1** | DMSO or FX11 | Lactate dehydrogenase A inhibitor solubilized in DMSO. | Test compound |
| **Injection 2** | Oligomycin mixture | Inhibitor of ATP synthase V of ETC | A decrease in OCR correlates cellular ATP generation to with mitochondrial respiration. |
| **Injection 3** | Carbonyl cyanide-4 (trifluoromethoxy) phenylhydrazone (FCCP) | Uncoupling agent results in maximum oxygen consumption rate by collapsing inner mitochondrial membrane. | An increase in OCR levels indicates maximum respiration capacity of cell. |
| **Injection 4** | Rotenone and antimycin A | Inhibitor of complexes I and III of ETC | A decrease in OCR correlates with shut down of mitochondrial respiration. Cellular respiration is driven by non-mitochondrial process. |

#### Glycolytic stress assay

Glucose is converted to pyruvate, and subsequently to lactate, results in proton generation and extrusion that acidify the extracellular medium (recorded as ECAR). This test was carried out to determine the impact of FX11 on ECAR values of BMDMs when sequentially treated with different glycolytic modulators.

|  | Glycolytic stress |  |  |
| --- | --- | --- | --- |
|  | compound | function | effect on ECAR |
| **Basal acidification** | Not applicable | - | Base line reading to assess non-glycolytic acidification |
| **Injection 1** | DMSO or FX11 | Lactate dehydrogenase A inhibitor solubilized in DMSO. | Test compound |
| **Injection 2** | Glucose | Glycolytic substrate. Glucose catabolism result in pyruvate and lactate and subsequent extracellular release of protons | An increase in ECAR value correlates with rate of glycolysis |
| **Injection 3** | Oligomycin mixture | Inhibitor of mitochondrial ATP synthase. Upon inhibition, cells are increasingly dependent on glycolysis. | Further increase in ECAR value correlates with maximum glycolytic capacity of the cell (in the absence of oxidative phosphorylation). |
| **Injection 4** | 2-deoxyglucose (2-DG) | Inhibitor of glucose hexokinase which mediates first step of glycolysis | A decrease in ECAR value implies that the ECAR produced in the experiment is due to glycolysis. |

## TEXT S2: Linear regression modelling analysis to determine the effect of FX11 on bone marrow derived macrophages bioenergetics and glycolytic response

### Data acquisition

Oxygen consumption (OCR) and extracellular acidification rates (ECAR) were measured using the Seahorse XF96 extracellular flux analyzer (Agilent, Santa Clara, CA). Two different assays were performed using the XF96: mitochondrial respiration assay, and glycolytic stress assay. Acquired real-time data were into the XF Report Generators using the Wave Desktop 2.6 software for calculation of the parameters from the specific assays.

## Results

### 1. Respiratory profile and respiratory parameters of BMDMs treated with FX11 or DMSO (vehicle control)

#### 1.1. Box plots showing respiratory response (OCR value) stratified by experiment replicate (related to Fig. 1A and B)

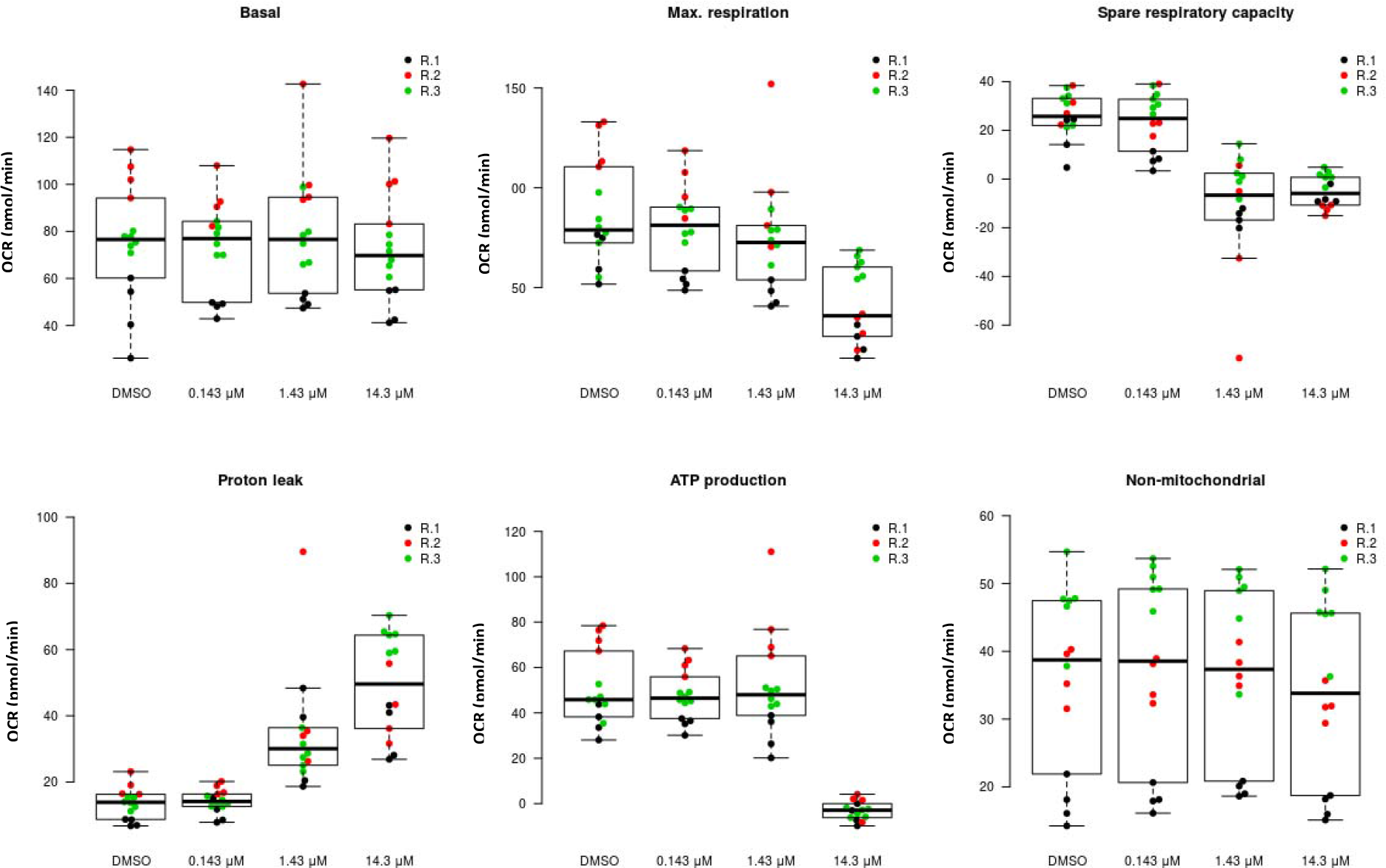

#### 1.2. Linear regression models for each of the six parameters (see above box plots in 1.1)

For each parameter (readout), the influence of FX11 concentration on the parameter readout was tested using log-linear regression. To this end, the FX11 concentrations were logarithmized (with the control, DMSO, assumed to have a concentration below 0.0143 mM) and a linear model (lm) was fit on the resulting data with the lm() function in R.

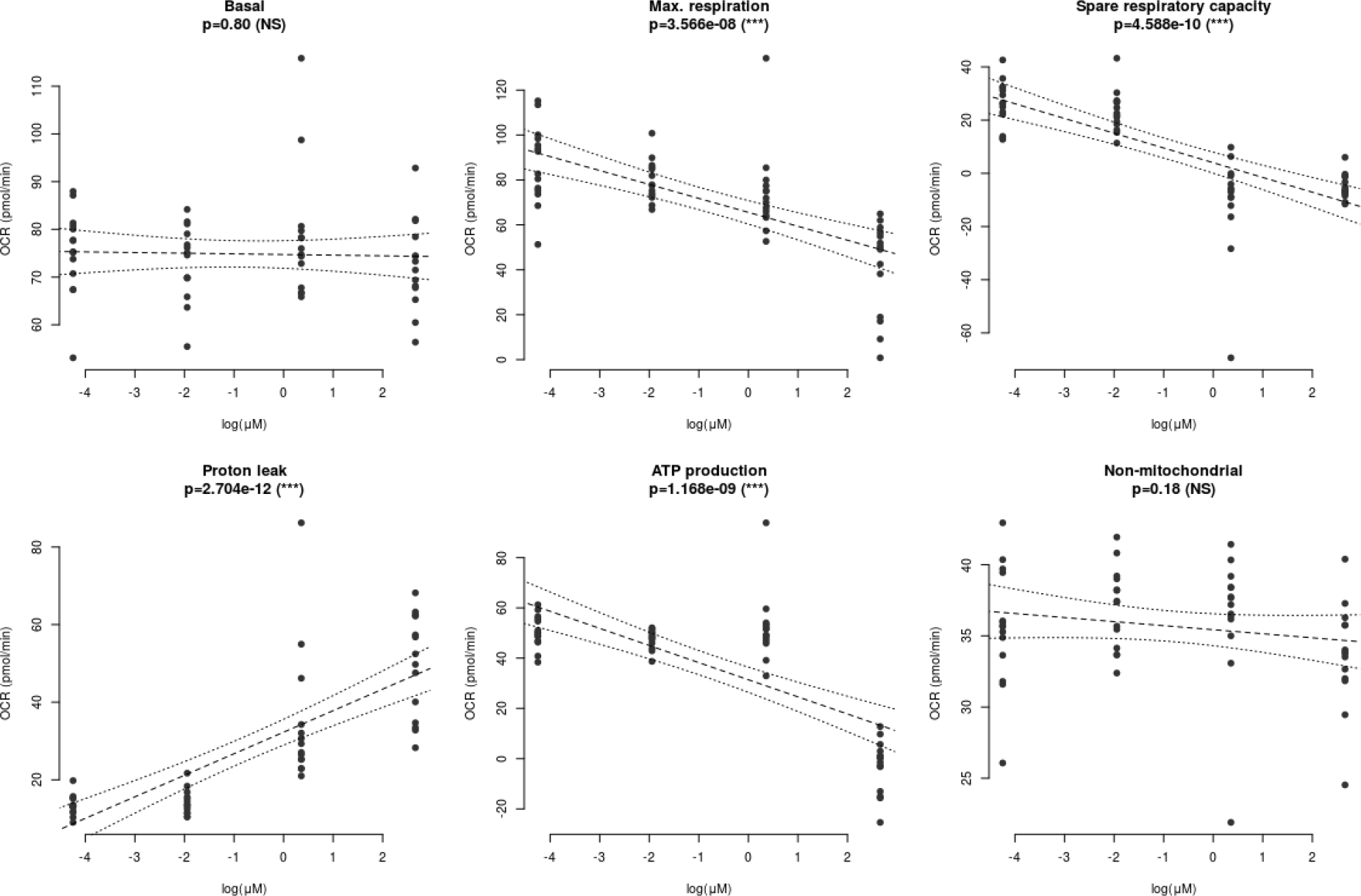

#### 1.3. Linear regression modeling results (related to respiratory parameters presented in Fig. 1A and B)

**1.3. Linear regression modeling results (related to respiratory parameters presented in Fig. 1A and B).**
| Parameter | P values |
| --- | --- |
| Basal | 0.80 (NS) |
| Max. respiration | 6.403e-08 (***) |
| Spare respiratory capacity | 9.652e-10 (***) |
| Proton leak | 6.864e-12 (***) |
| ATP production | 2.375e-09 (***) |
| Non-mitochondrial | 0.18 (NS) |

### 2. Glycolytic stress profile and glycolytic parameters of BMDMs-treated with FX11 or DMSO (vehicle control)

#### 2.1. Box plots showing glycolytic response (ECAR value) stratified by experiment replicate (related to Fig. 1C and D)

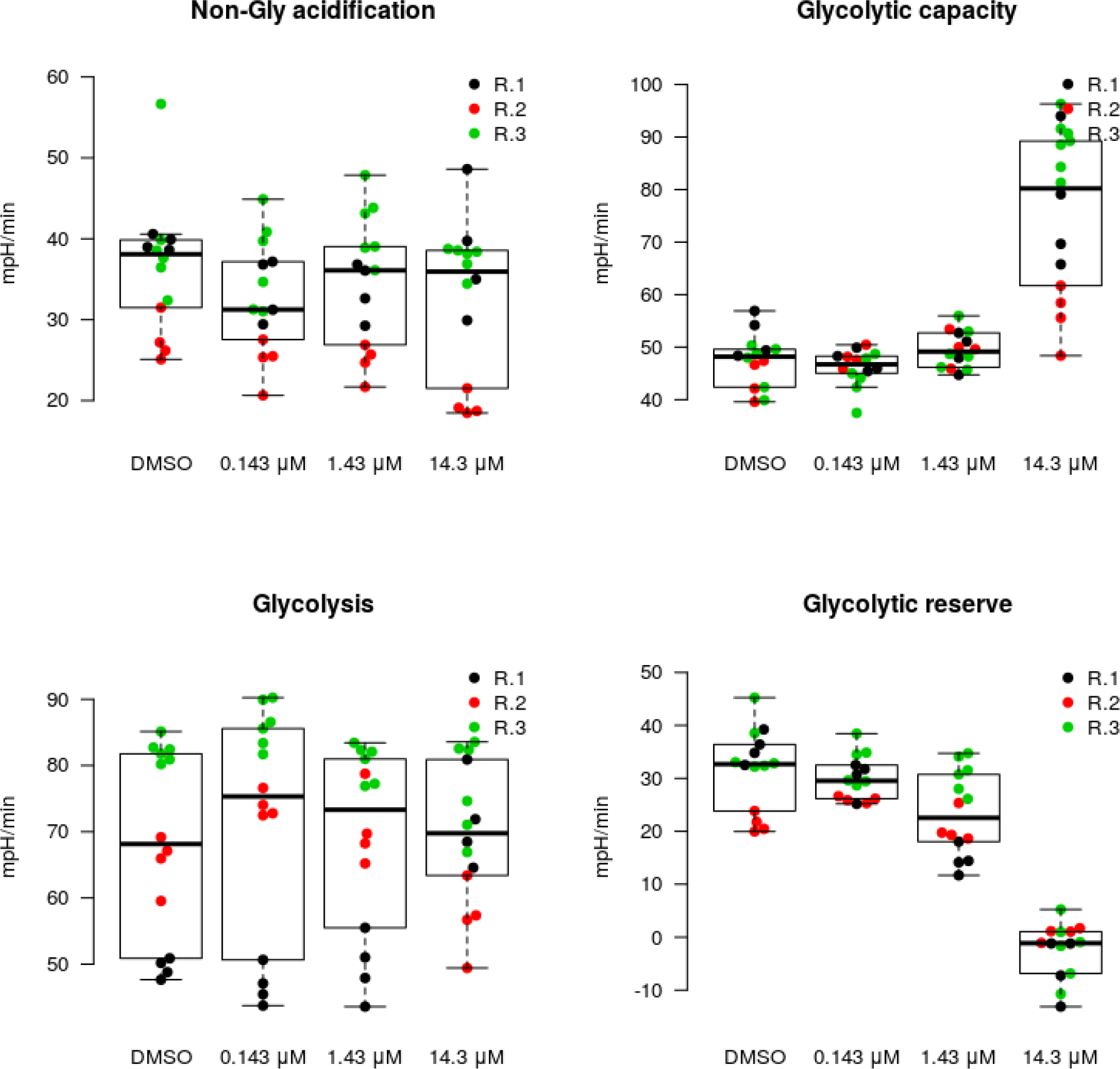

#### 2.2. Linear regression models for each of the four outputs collected (see above box plots in 2.1.)

For each parameter (readout), the influence of FX11 concentration on the parameter readout was tested using log-linear regression. To this end, the FX11 concentrations were logarithmized (with the control, DMSO, assumed to have a concentration below 0.0143 mM) and a linear model (lm) was fit on the resulting data with the lm() function in R.

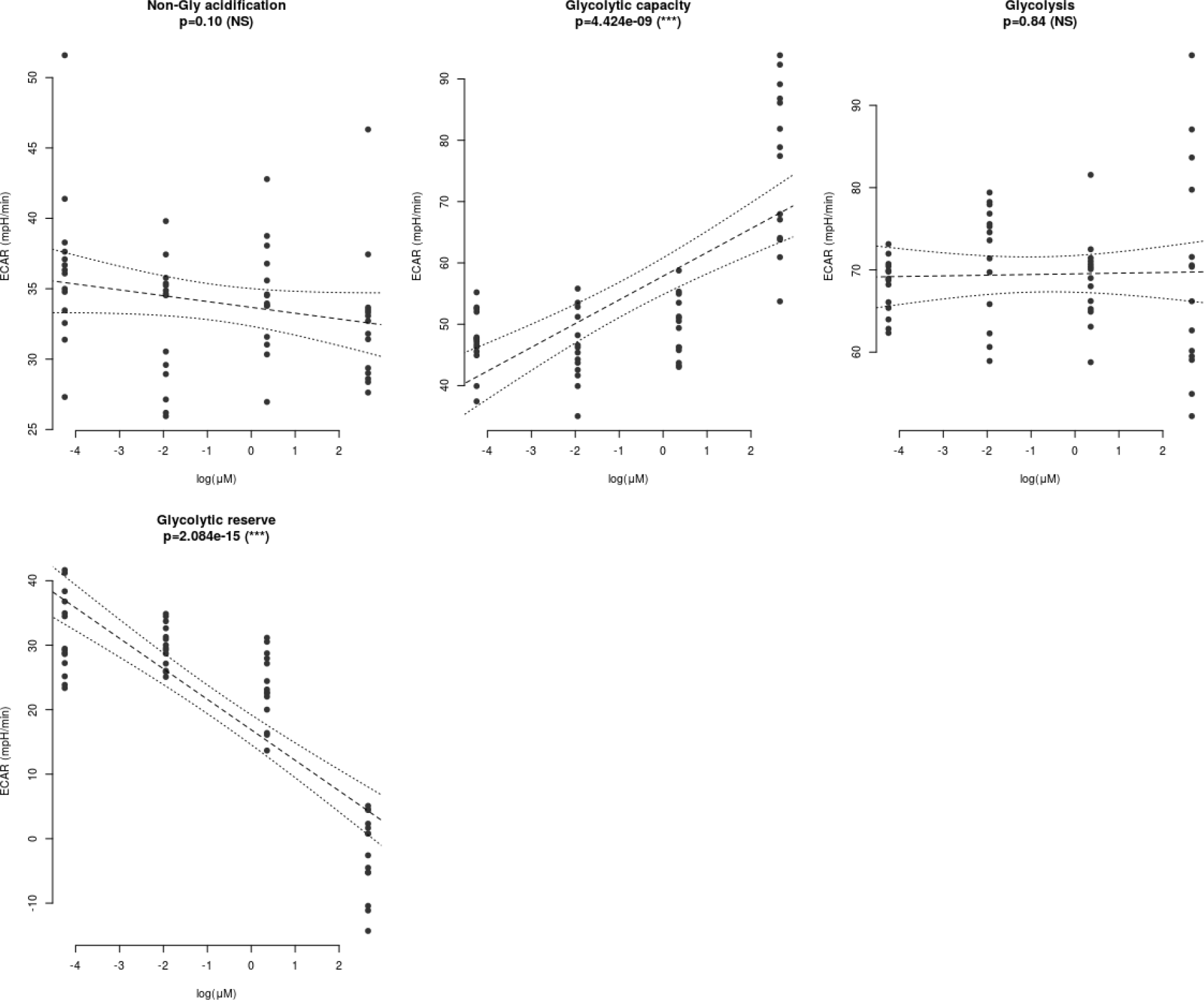

#### 2.3 Linear regression modeling results (related to glycolytic function parameters presented in Fig. 1C and D)

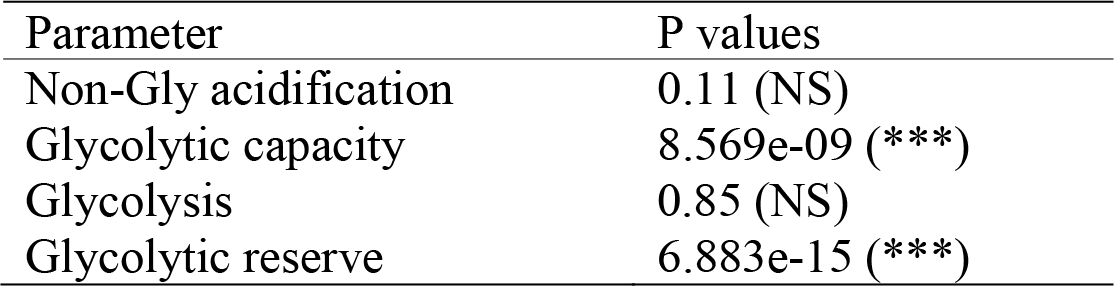

**FIG S1.**
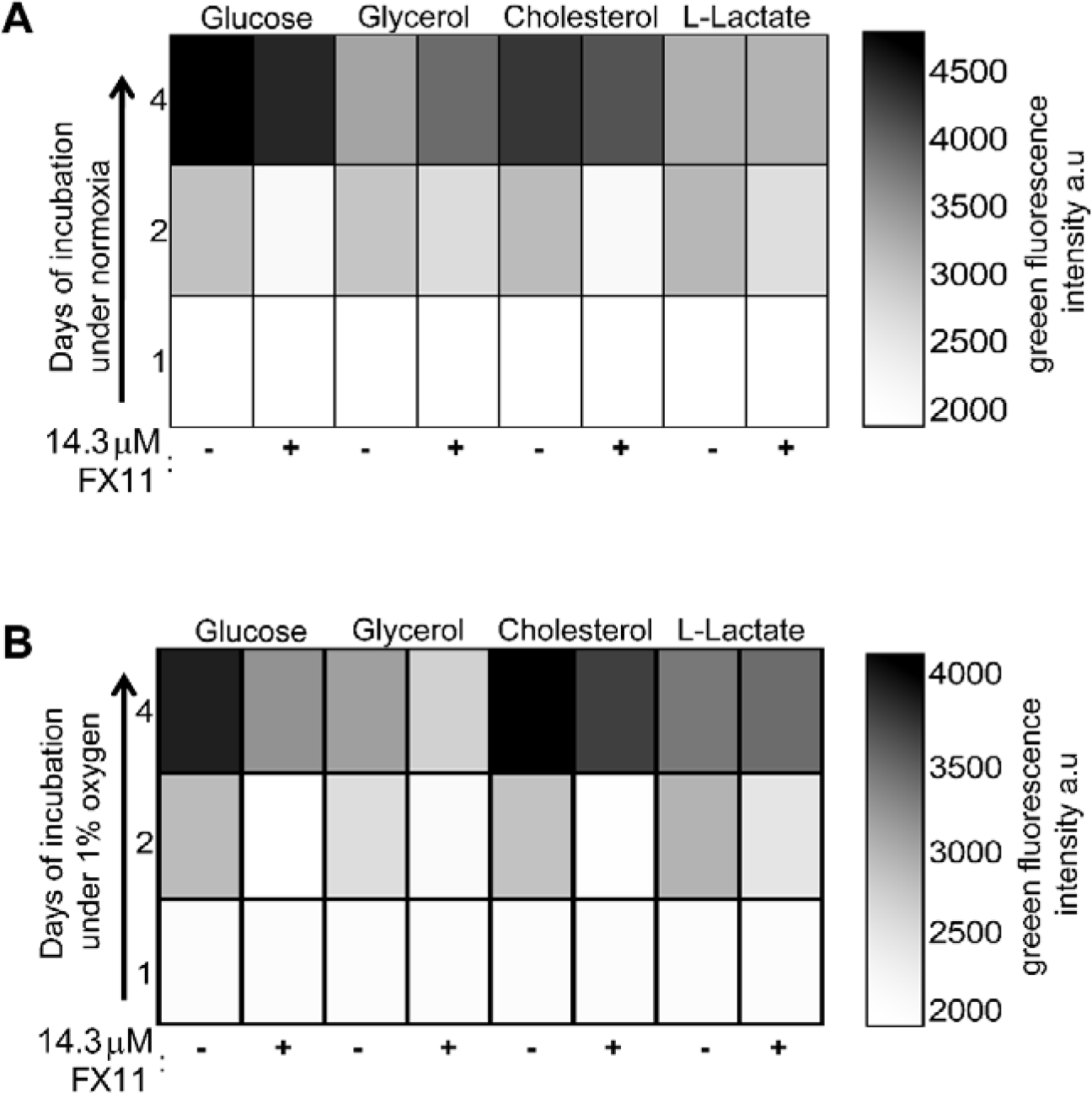
Effects of FX11 on bacterial growth. **(A-B)** Gradient color map showing the fluorescence intensity of green fluorescence protein expressing *M. tuberculosis* strain. Liquid culture in medium containing specified carbon sources and incubated under aerobic or hypoxic growth condition at 37° C.

**FIG S2.**
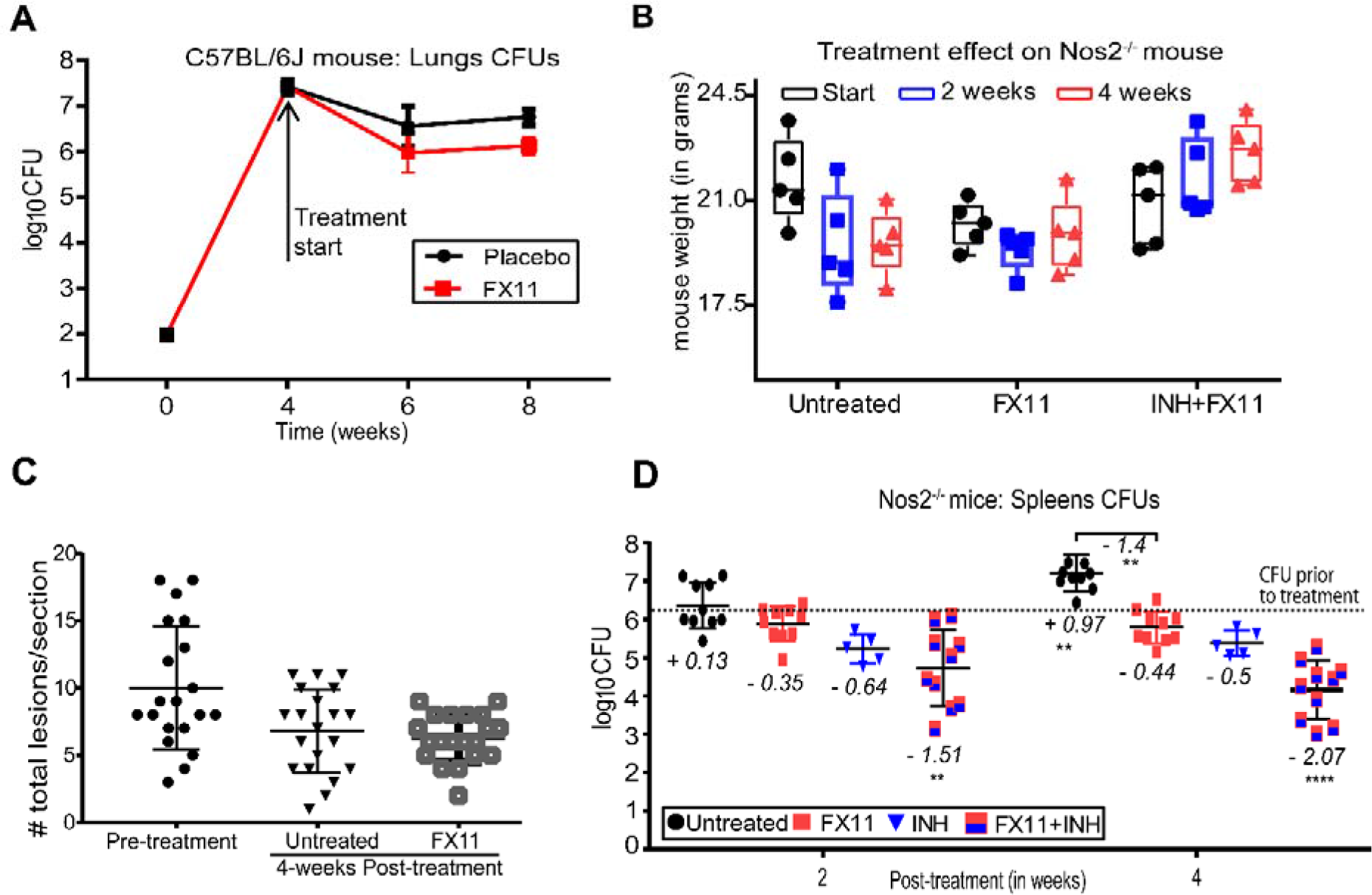
Effects of FX11 administration into mouse models. **(A)** Bacterial burden in C57BL/6J lungs are shown at respective time points. **(B)** Body weight of untreated and drug-treated Nos2^−/−^ mice. **(C)** Total number of lesions (necrotic and non-necrotic) per lung section of Nos2^−/−^ mouse groups. **(D)** Splenic CFU data (means±SD) from two independent experiments (n = 9–10). Italicized numerical value represents log_10_CFU differences (increase in CFU is indicated as positive value and decrease is indicated as negative value) of the specified group, when compared with the control group prior to drug treatment (i.e. day 56, indicated in dotted line). Pooled data from two independent experiments were analyzed using nonparametric Mann-Whitney test (data that did not pass the Shapiro-Wilk normality test). Statistical significance as compared to the group prior to drug treatment, *p<0.05, **p<0.01, ****p<0.0001.

**FIG S3.**
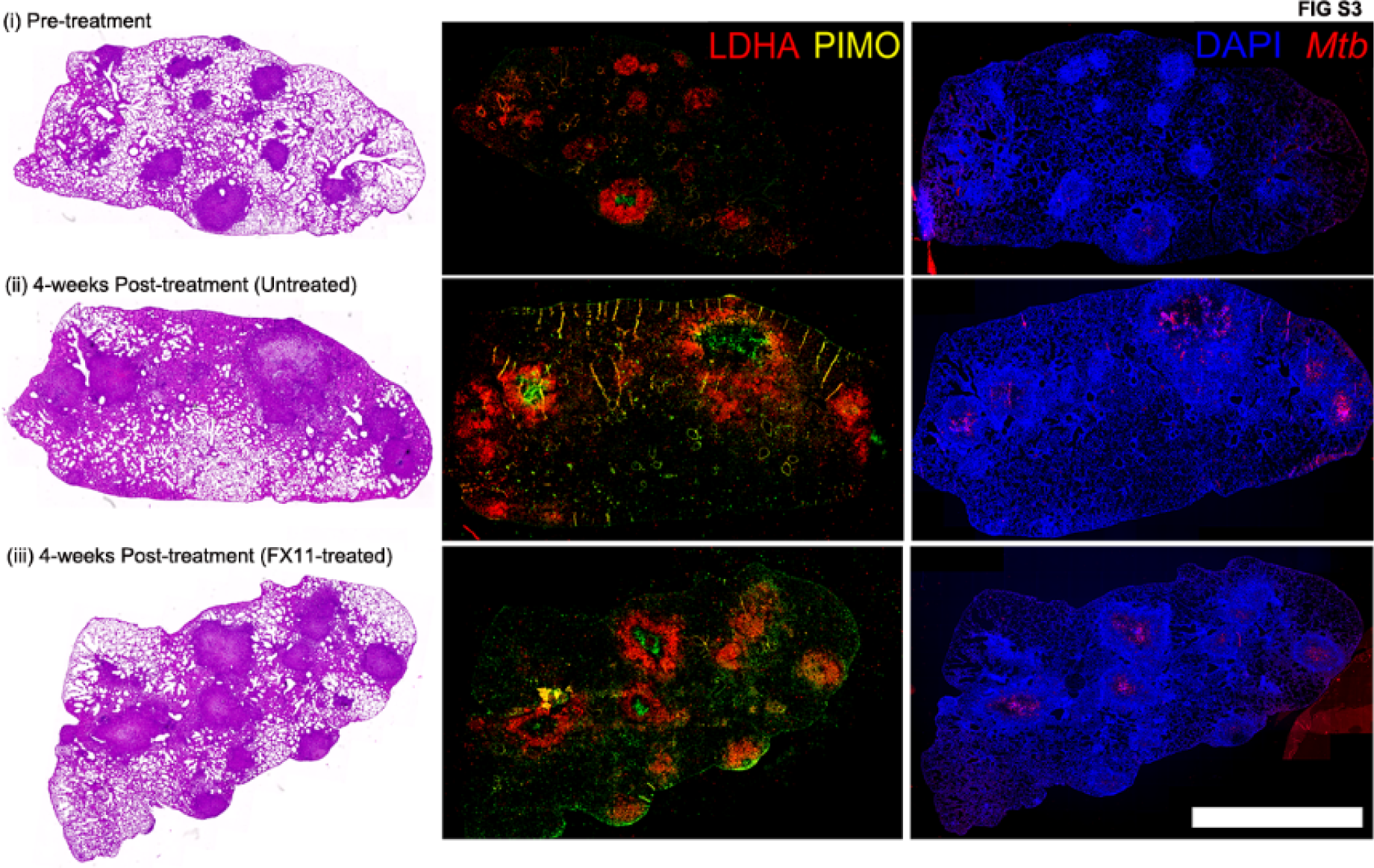
Staining of whole lung section from Nos2^−/−^ mice. Micrographs of stained consecutive thin sections of the fixed and paraffin-embedded left lung lobe. Scale bar represents 2.5 mm.

